# 53BP1 mediates sensitivity to chemotherapy and is associated with poor clinical outcomes in high-grade serous ovarian cancer

**DOI:** 10.1101/2023.09.30.560286

**Authors:** Michael Skulimowski, Jessica Bourbonnais, Nicolas Malaquin, Hubert Fleury, Isabelle Clément, Laudine Communal, Kurosh Rahimi, Diane Provencher, Anne-Marie Mes-Masson, Francis Rodier

## Abstract

High-grade serous ovarian cancer (HGSOC) remains the most lethal gynecological malignancy in North American women. At a cellular level, the current first-line chemotherapies cause DNA-damage and activate the DNA damage response signalling cascade. Here we explore the role of 53BP1, a central mediator of the DNA damage response, in HGSOC chemotherapy outcomes. Tissue 53BP1 protein levels were quantified in two independent HGSOC cohorts, the COEUR validation cohort (n = 173) and CHUM cohort (n = 56). Univariate and multivariate analyses showed that high nuclear 53BP1 levels in ovarian cancer cells were strongly associated with poor disease-specific survival in both cohorts. High 53BP1 was associated with poor progression-free survival (PFS) in the COEUR cohort, and trended towards poor PFS in the CHUM cohort. These findings were validated by whole-tumour *TP53BP1* mRNA of the TCGA Firehose Legacy cohort (n = 591) in which high *TP53BP1* mRNA levels were associated with poor overall survival on multivariate analysis. In HGSOC cell lines, 53BP1 levels were positively correlated with resistance to carboplatin using colony formation assay, and depletion of 53BP1 sensitized resistant cell lines to genotoxic therapies. These results suggest that 53BP1 is associated with poor prognosis in HGSOC and may mediate this relationship by modulating cellular sensitivity to chemotherapy.

**Statement of translational relevance:** Current first-line chemotherapies in ovarian cancer cause DNA damage and activate the DNA damage response, culminating in the taking of cell fate decisions. 53BP1 is a central mediator in this signalling cascade, where it is involved at multiple levels: signal amplification, recruitment of effectors, DNA repair pathway choice, and cell cycle regulation. However, its role in ovarian cancer treatment outcomes remains unknown. In this study, we found that 53BP1 correlated with poor clinical outcomes in three ovarian cancer patient cohorts and mediated carboplatin sensitivity in ovarian cancer cells. These results reveal 53BP1 and the DNA damage response as important actors in ovarian cancer treatment response. Though further studies are necessary to gain a more complete understanding of their involvement in clinical outcomes, they appear as promising candidates for potential therapeutic targeting in ovarian cancer.

## INTRODUCTION

Ovarian cancer is the second most common and the most lethal gynecological malignancy in North America (Siegel et al., 2020). High-grade serous ovarian cancer (HGSOC) is by far the most frequent histological subtype, making up nearly 70 % of cases (Prat, 2012). Current first line therapies consist of a combination of platinum- and taxane-based chemotherapies and surgical cytoreduction (Lheureux et al., 2019; Network, 2020). Although initial response rates can be as high as 80 %, only 20 % of patients remain disease-free at 5 years (Lheureux et al., 2019). Recurrence or persistence of disease are invariably associated with a terminal clinical course, eventually culminating in the patient’s death after cycles of response and relapse (Lheureux et al., 2019). With only modest improvements of survival rates in ovarian cancer over the past 20 years (Committee, 2019), these poor clinical outcomes highlight the need for a better fundamental understanding of treatment responses in HGSOC.

At the molecular level, current HGSOC treatments induce DNA damage, including DNA double-stranded breaks (DSBs) (Sorenson and Eastman, 1988; Wang et al., 2020), through a variety of mechanisms (Basu and Krishnamurthy, 2010; Weaver, 2014). This triggers the DNA damage response (DDR) signalling cascade, which leads to the activation of cell cycle checkpoints and DNA repair pathways, and eventually culminates in the taking of cell fate decisions, including, but not limited to, a return to the cell cycle and continued proliferation, or cell death via cellular senescence, apoptosis or mitotic catastrophe (d’Adda di Fagagna, 2008; Minafra and Bravatà, 2014). By mediating outcomes at a cellular level, the DDR mediates outcomes at the clinical level as well. Strikingly, HGSOC is characterized by mutations at multiple points in the DDR, sometimes providing novel targets to enhance therapy. *TP53* encodes p53, one of the main downstream effectors of the DDR involved in treatment-mediated cell fate decisions, and is almost ubiquitously dysfunctional in this disease (Cancer Genome Atlas Research, 2011). DNA repair pathways are also altered, with nearly half of patients harbouring defects in the homologous recombination (HR) DSB repair pathway (Cancer Genome Atlas Research, 2011), providing targets for emerging DNA damaging therapies involving poly (ADP-ribose) polymerase (PARP) inhibitors (Fleury et al., 2019; Ledermann et al., 2012) . As we have recently shown, combining DNA damage repair and vulnerabilities in the cell fate decisions in HGSOC can yield novel approaches to treat this disease (Fleury et al., 2019). Overall, this renders the DDR an exceptionally interesting pathway to study in the context of HGSOC.

Chief among the various actors in the DDR is 53BP1 (*TP53BP1*). The most well-known role of 53BP1 in this signalling cascade is to direct DNA repair pathway choice. By protecting DSB ends from resection and by counteracting BRCA1, 53BP1 promotes non-homologous end joining (NHEJ) and inhibits HR (Scully et al., 2019). Most notably, the promotion of aberrant NHEJ by 53BP1 in BRCA1-deficient (i.e., HR-deficient) cells is in part responsible for the toxicity of PARP inhibitors (Bunting et al., 2010), a drug class now used in the treatment of breast and ovarian cancers (Tew et al., 2020; Tung et al., 2020). Accordingly, downregulation of 53BP1 in mouse models has been suggested as a mechanism of resistance to PARP inhibitors (Jaspers et al., 2013). However, 53BP1 plays multiple additional roles in the DDR. For instance, 53BP1 is recruited to the site of DSBs where it acts as a scaffold for the recruitment of several other DDR factors, including chromatin modulator EXPAND1 (Huen et al., 2010), RIF1 (Silverman et al., 2004), and PTIP (Munoz et al., 2007) (Noordermeer et al., 2018). Despite this, 53BP1 knockdown has only a minimal effect on the kinetics of DSB repair (Noon et al., 2010).

53BP1 also plays a role in checkpoint reinforcement in the S and G2-M phases of the cell cycle by promoting phosphorylation of multiple ATM substrates, including checkpoint kinase 2 (CHK2) (Batenburg et al., 2017; DiTullio et al., 2002; Fernandez-Capetillo et al., 2002; Ward et al., 2003). Evidence suggests this maintenance of cell cycle checkpoints by 53BP1 may help promote genomic stability, particularly in p53-deficient cells (Morales et al., 2006). Lastly, 53BP1 directly interacts with p53, potentially modulating cell fate decisions downstream of the DDR. However, this function remains poorly understood (Brummelkamp et al., 2006; Cuella-Martin et al., 2016).

Given its involvement at multiple levels in the DDR, the role of 53BP1 in cancer and cancer treatment is most likely dependent on the integrity of different pathways that interact with 53BP1, as evidenced by the heterogeneity of effects of 53BP1 on cellular responses to treatment reported in the literature. Among the very limited literature on 53BP1 in ovarian cancer, Pennington et al. found that lower expression of 53BP1 mRNA in tissue was associated with improved survival in the absence of *BRCA1/2* mutations in a cohort including primary or recurrent epithelial ovarian, fallopian tube, and peritoneal carcinoma patients (Pennington et al., 2013). However, they reported no correlation between 53BP1 immunohistochemical staining intensity and clinical outcomes (Pennington et al., 2013). Alternatively, Hurley et al. showed that reducing 53BP1 levels in a BRCA1-deficient ovarian cancer cell line increased HR and decreased sensitivity to PARP inhibitors, consistent with their observation that the low expression of 53BP1 in ovarian cancer patients with HR-deficient tumours correlated with decreased short-term response to PARP inhibitors (Hurley et al., 2019). Thus, while important roles are suggested and possibly contrasted, much about 53BP1 in HGSOC remains undefined.

In this study we focus on using HGSOC patient tissues and their associated clinical data, and HGSOC cell lines, we show that high levels of nuclear 53BP1 from quantitative immunofluorescence (IF) on treatment-naïve HGSOC tissues correlated with poor disease-specific survival in univariate and multivariate analyses on two independent cohorts, and similar trends were observed with *TP53BP1* mRNA levels in a third cohort. We also show that 53BP1 expression in treatment-naïve HGSOC cell lines correlated with carboplatin sensitivity, and the depletion of 53BP1 sensitized cell lines to carboplatin. Our findings suggest that 53BP1 plays a key role in HGSOC treatment outcomes and warrants further investigation.

## RESULTS

### Demographic characteristics of the COEUR and CHUM cohorts

To explore the role of 53BP1 in HGSOC treatment, we first correlated 53BP1 protein levels in HGSOC tissues with clinical outcomes. We obtained tissue specimens in the form of tissue microarrays (TMAs) from two Canadian cohorts, along with associated clinical data. The COEUR validation cohort included tissues from 173 HGSOC patients recruited between 1992 and 2011 at various Canadian institutions, including the CHUM. The CHUM cohort was an in-house cohort of 76 HGSOC patients recruited between 1993 and 2012, which was used to validate results obtained from the COEUR cohort. Twenty patients belonged to both the CHUM and COEUR validation cohorts and were removed from analysis in the CHUM cohort (final n=56). Only patients naïve to chemotherapy were included in the study to avoid any selection bias posed by post-chemotherapy tissue collection. The clinical characteristics of the patients included in each cohort are described in Table 1.

**Table 1:** Clinical characteristics of the patients in the COEUR and CHUM cohorts.

| Characteristics | COEUR (n = 173) | CHUM (n = 56) |
| --- | --- | --- |
| Age at diagnosis – years |  |  |
| Median | 63.0 | 60.0 |
| Range | 34.3-89.0 | 34.0-79.0 |
| FIGO stage at presentation – n (%) |  |  |
| 1 | 19 (11.0) | 2 (3.6) |
| 2 | 26 (15.0) | 4 (7.1) |
| 3 | 116 (67.1) | 43 (76.8) |
| 4 | 11 (6.4) | 7 (12.5) |
| Unknown | 1 (0.6) | 0 (0) |
| Residual disease after surgery – n (%) |  |  |
| None | 45 (26.0) | 11 (19.6) |
| 1 cm or less | 43 (24.9) | 12 (21.4) |
| 1-2 cm | 19 (11.0) | 9 (16.1) |
| Greater than 2 cm | 45 (26.0) | 18 (32.1) |
| Milliary | 3 (1.7) | 2 (3.6) |
| Unknown | 18 (10.4) | 4 (7.1) |
| BRCA-status – n (%) |  |  |
| BRCA1-mutated | 12 (6.9) | 2 (3.6) |
| BRCA2-mutated | 8 (4.6) | 2 (3.6) |
| Wild-type | 105 (60.7) | 44 (78.6) |
| Unknown | 48 (27.7) | 8 (14.3) |
| First-line chemotherapy – n (%) |  |  |
| Carboplatin and paclitaxel | 127 (73.4) | 21 (37.5) |
| Cisplatin and paclitaxel | 20 (11.6) | 5 (8.9) |
| Carboplatin | 6 (3.5) | 0 (0.0) |
| Cisplatin | 5 (2.9) | 1 (1.8) |
| Other | 12 (6.9) | 9 (16.1) |
| None | 1 (0.6) | 1 (1.8) |
| Unknown | 2 (1.2) | 19 (33.9) |
| Follow up – months |  |  |
| Median | 48.3 | 38.0 |
| Range | 0.2-185.0 | 3.0-156.0 |
| Disease-specific survival status – n (%) |  |  |
| Alive | 85 (49.1) | 19 (33.9) |
| Deceased | 86 (49.7) | 31 (55.4) |
| Unknown | 2 (1.2) | 6 (10.7) |
| Disease-specific survival time – months |  |  |
| Median | 91.3 | 53.0 |
| Range | 0.4-185.0 | 3.0-156.0 |
| Progression-free survival status – n (%) |  |  |
| No progression | 71 (41.0) | 8 (14.3) |
| Progression | 99 (57.2) | 47 (83.9) |
| Unknown | 3 (1.7) | 1 (1.8) |
| Progression-free survival time – months |  |  |
| Median | 24.0 | 16.0 |
| Range | 0.4-185.0 | 1.0-97.0 |
| Tissue year of collection |  |  |
| Median | 2005 | 2003 |
| Range | 1992-2011 | 1993-2012 |

### High 53BP1 protein levels in HGSOC tissues correlate with poor clinical outcomes in univariate analysis

We quantified 53BP1 protein levels in HGSOC tissues using quantitative multicolor IF. TMAs from each cohort were stained with DAPI for identification of nuclei, as well as for cytokeratins 7, 18 and 19 for identification of the epithelial (i.e., tumoral) compartment, and for 53BP1 (Fig. 1a). IF images were then digitalized at high resolution and quantitatively analyzed using computer-assisted segmentation/analysis. The epithelial compartment and nuclei were identified in each core using the cytokeratin mask signal and DAPI signal, respectively. Each core was separated into epithelial and stromal compartments, and each compartment was subdivided into nuclear and cytoplasmic compartments (Fig. 1b). The mean fluorescent intensity (MFI) of the 53BP1 signal was then quantified in the epithelial nuclear compartment of each core (i.e. in cancer cell nuclei). The average MFI across two duplicate cores was calculated for each patient and subsequently used for clinical correlations. Separation of patients into high and low expressors of 53BP1 was carried out by determining optimal cut-off points on the receiver operating characteristic curves of 53BP1 MFI in relation to clinical outcomes (disease-specific survival and progression-free survival). Cut-offs were determined for each outcome in each cohort.

**Figure 1:**
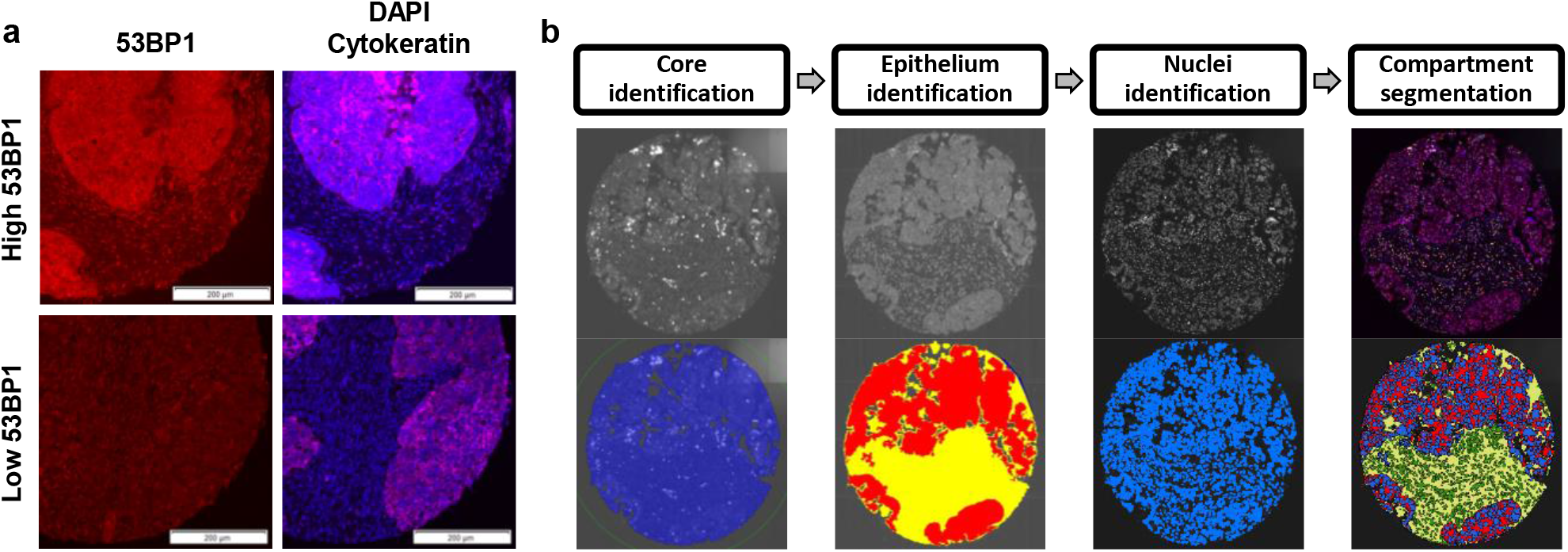
Computer-assisted image segmentation and analysis of immunofluorescence-stained HGSOC tissue cores. (**a**) Representative images of tissue cores staining strongly (top two panels) and weakly (bottom two panels) for 53BP1 immunofluorescence. Left: 53BP1 (red). Right: DAPI (blue); cytokeratin 7, 18 and 19 (fuscia). (**b**) Computer-assisted image analysis workflow. TMA cores were simultaneously stained for 53BP1, DAPI, and cytokeratins 7, 18 and 19, each in a separate channel. We first identified cores from the background using a fluorescence threshold (far left). The epithelial (i.e., tumoral) and stromal compartments were identified using the cytokeratin mask and a fluorescence threshold, and saved as regions of interest (ROIs) (middle left). Nuclei were subsequently identified using a fluorescence threshold on the DAPI staining (middle right) and saved as ROIs. The tissue cores were then subdivided into epithelial and stromal, as well as nuclear and cytoplasmic compartments using the ROIs previously identified (far right). The MFI of 53BP1 was thereafter analyzed in the epithelial nuclear compartment.

Using Kaplan-Meier survival plots to determine disease-specific survival and progression-free survival, we compared the high and low 53BP1 groups in each cohort. In the COEUR validation cohort, high 53BP1 expression was associated with strikingly poorer disease-specific survival (log-rank, p < 0.001): low expressors had a mean disease-specific survival of 106.9 months (95% confidence interval (CI): 91.5-122.2), whereas high expressors had a mean of 73.7 months (95% CI: 59.9-87.5) (Fig. 2a). Accordingly, patients with high epithelial nuclear 53BP1 were found to have significantly poorer progression-free survival (log-rank, p < 0.001) with a mean progression-free survival of 46.0 months (95% CI: 31.8-60.2) as compared to the 93.2 months (95% CI: 78.1-108.3) of the low 53BP1 group (Fig. 2a). We sought next to validate these results with our in-house CHUM cohort. Remarkably, high epithelial nuclear 53BP1 was associated with significantly poorer disease-specific survival in the CHUM cohort (log-rank, p = 0.0325) (Fig. 2b) as observed in the COEUR validation cohort. Patients with high 53BP1 had a mean disease-specific survival of 49.2 months (95% CI: 34.9-63.6), whereas patients with low 53BP1 had a mean disease-specific survival of 90.8 months (95% CI: 57.2-124.4). Although statistical significance was not achieved, the high 53BP1 group showed a strong trend for decreased progression-free survival as compared to the low 53BP1 group (log-rank, p = 0.0536) (Fig. 2b). Progression-free survival was 10.4 months (95% CI: 5.2-15.6) in the high 53BP1 group and 29.4 months (95% CI: 21.0-37.7) in the low 53BP1 group. Thus, in univariate Kaplan-Meier analysis, high epithelial nuclear 53BP1 levels in HGSOC tissues were associated with poorer disease-specific survival and progression-free survival in two independent cohorts.

**Figure 2:**
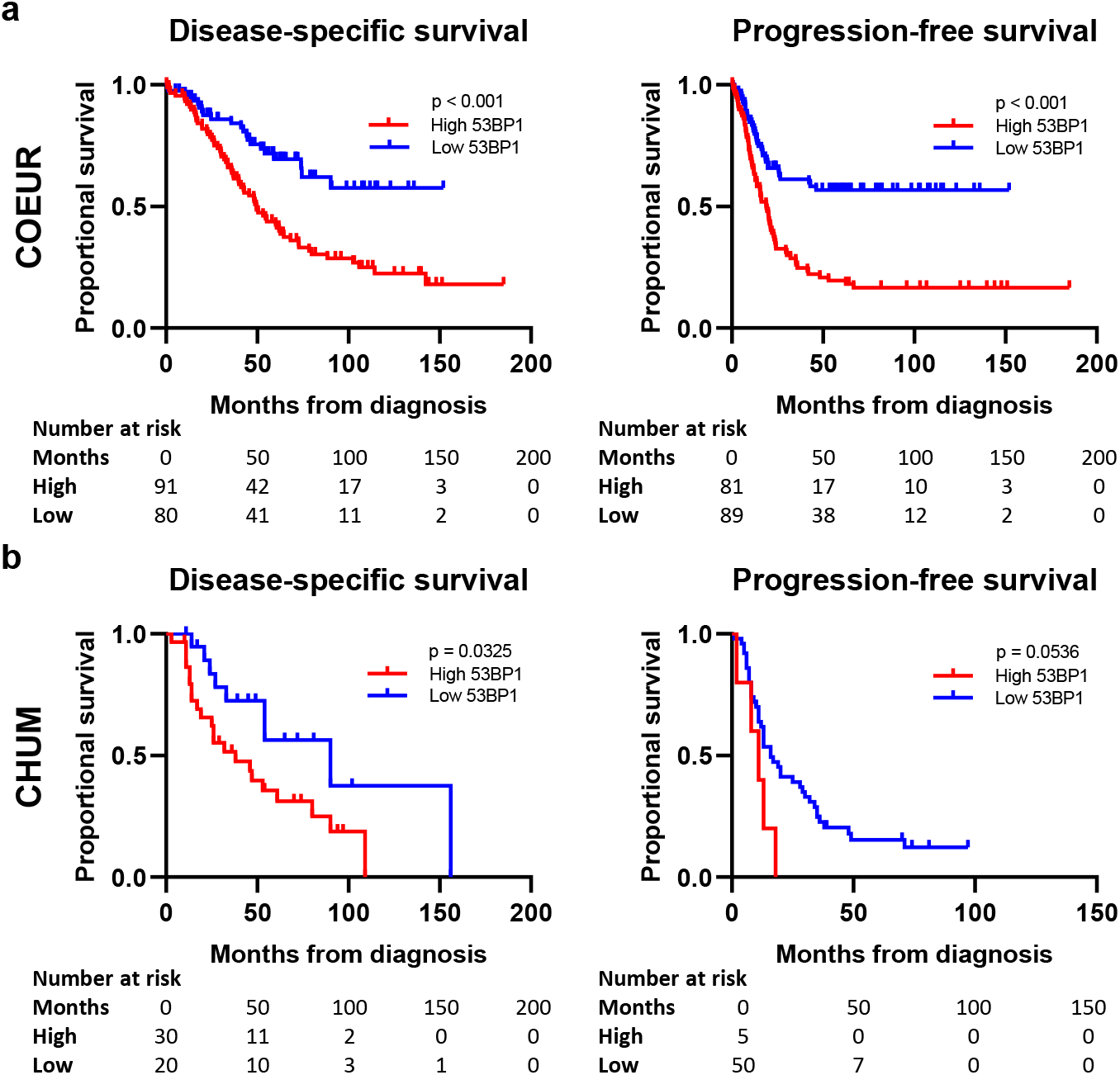
53BP1 is associated with disease-specific survival and progression-free survival in the COEUR and CHUM cohorts. Kaplan-Meier curves for disease-specific survival (left) and progression-free survival (right) for high and low expressors of epithelial nuclear 53BP1 in the **(a)** COEUR cohort and (**b**) CHUM cohort. Statistical significance was calculated using the log-rank test.

### High 53BP1 protein levels in HGSOC tissues correlate with poor clinical outcomes in multivariate analysis

We next investigated whether epithelial nuclear 53BP1 is predictive of clinical outcomes in HGSOC, independent of other prognostic factors. We performed univariate and multivariate Cox regression analyses with 53BP1 as a dichotomous (high vs. low) variable per the cut-offs previously established, and as a continuous variable for disease-specific survival and progression-free survival. Three prognostic clinical variables including age at diagnosis, disease stage at presentation, and residual disease after surgery were controlled in the multivariate analysis. Disease stage at presentation and residual disease after surgery were significantly associated with both outcomes studied in the COEUR validation cohort (Table 2a). In the CHUM cohort, residual disease after surgery was associated with poorer progression-free survival and trended towards poorer disease-specific survival, while disease stage at presentation trended towards a poorer prognosis with both outcomes (Table 2b). Age at diagnosis was not associated with either outcome in either cohort.

**Table 2:** Cox regression analysis of 53BP1 expression in the COEUR and CHUM cohorts. Cox proportional hazard regression analysis of selected variables and disease-specific (left) or progression-free (right) survival in the **(a)** COEUR cohort and **(b)** CHUM cohort. Age at diagnosis and nuclear epithelial 53BP1 expression (where stated) were continuous variables. ^a^Hazard ratio (HR) estimated from the Cox proportional hazard regression model. ^b^95% confidence interval around the estimated HR. ^c^Multivariate analysis was adjusted for three clinical variables: age at diagnosis, disease stage at presentation, and residual disease after surgery. Statistically significant p-values in bold.

| Variable | COEUR |  |  |  |  |  |
| --- | --- | --- | --- | --- | --- | --- |
|  | Disease-specific survival |  |  | Progression-free survival |  |  |
|  | HR<sup>a</sup> | 95 % CI<sup>b</sup> | p-value | HR<sup>a</sup> | 95 % CI<sup>b</sup> | p-value |
| Age at diagnosis | 0.999 | 0.979-1.019 | 0.912 | 0.993 | 0.975-1.012 | 0.467 |
| Disease stage |  |  | **5.72x10<sup>-3</sup>** |  |  | **2.9x10<sup>-5</sup>** |
| 2 vs 1 | 0.753 | 0.230-2.470 | 0.640 | 0.743 | 0.227-2.435 | 0.624 |
| 3 vs 1 | 2.417 | 0.973-6.000 | 0.0572 | 3.503 | 1.417-8.661 | **6.64x10<sup>-3</sup>** |
| 4 vs 1 | 3.786 | 1.197-11.98 | **0.0235** | 6.358 | 2.070-19.53 | **1.24x10<sup>-3</sup>** |
| Residual disease after surgery |  |  | **3.0x10<sup>-6</sup>** |  |  | **1.40x10<sup>-9</sup>** |
| 1 cm or less vs none | 2.085 | 0.949-4.582 | 0.0673 | 3.178 | 1.531-6.594 | **1.91x10<sup>-3</sup>** |
| 1-2 cm vs none | 2.485 | 1.008-6.121 | **0.0479** | 2.608 | 1.085-6.269 | **0.0321** |
| Greater than 2 cm vs none | 6.156 | 2.954-12.83 | **1.0x10<sup>-6</sup>** | 9.347 | 4.619-18.91 | **5.1x10<sup>-10</sup>** |
| Miliary vs none | 3.595 | 0.774-16.69 | 0.102 | 4.659 | 1.020-21.29 | **0.0471** |
| **53BP1 expression (continuous)** |  |  |  |  |  |  |
| Univariate | 1.007 | 1.002-1.013 | **0.0134** | 1.010 | 1.005-1.016 | **3.79x10<sup>-4</sup>** |
| Adjusted for clinical variables<sup>c</sup> | 1.007 | 1.001-1.014 | **0.0310** | 1.009 | 1.002-1.015 | **5.96x10<sup>-3</sup>** |
| **53BP1 expression (high vs. low)** |  |  |  |  |  |  |
| Univariate | 2.450 | 1.508-3.981 | **2.95x10<sup>-4</sup>** | 2.651 | 1.747-4.023 | **5.0x10<sup>-6</sup>** |
| Adjusted for clinical variables<sup>c</sup> | 2.325 | 1.305-4.141 | **4.18x10<sup>-3</sup>** | 2.179 | 1.349-3.518 | **1.45x10<sup>-3</sup>** |

| Variable | CHUM |  |  |  |  |  |
| --- | --- | --- | --- | --- | --- | --- |
|  | Disease-specific survival |  |  | Progression-free survival |  |  |
|  | HR<sup>a</sup> | 95 % CI<sup>b</sup> | p-value | HR<sup>a</sup> | 95 % CI<sup>b</sup> | p-value |
| Age at diagnosis | 0.995 | 0.954-1.037 | 0.799 | 0.994 | 0.962-1.027 | 0.713 |
| Disease stage |  |  | 0.235 |  |  | 0.133 |
| 2 vs 1 | 0.891 | 0.055-14.44 | 0.935 | 1.959 | 0.203-18.90 | 0.561 |
| 3 vs 1 | 1.486 | 0.199-11.09 | 0.699 | 2.776 | 0.378-20.40 | 0.316 |
| 4 vs 1 | 3.861 | 0.439-33.98 | 0.223 | 6.355 | 0.763-52.93 | 0.0873 |
| Residual disease after surgery |  |  | 0.0842 |  |  | **0.00977** |
| 1 cm or less vs none | 2.213 | 0.425-11.52 | 0.345 | 0.883 | 0.331-2.358 | 0.804 |
| 1-2 cm vs none | 3.925 | 0.760-20.27 | 0.103 | 1.728 | 0.646-4.626 | 0.276 |
| Greater than 2 cm vs none | 7.048 | 1.579-31.45 | **0.0105** | 3.575 | 1.529-8.358 | **0.00328** |
| Miliary vs none | 5.275 | 0.738-37.68 | 0.0974 | 2.956 | 0.603-14.48 | 0.181 |
| **53BP1 expression (continuous)** |  |  |  |  |  |  |
| Univariate | 1.002 | 0.999-1.005 | 0.160 | 1.001 | 0.999-1.003 | 0.408 |
| Adjusted for clinical variables<sup>c</sup> | 1.003 | 0.999-1.007 | 0.217 | 1.001 | 0.998-1.004 | 0.397 |
| **53BP1 expression (high vs. low)** |  |  |  |  |  |  |
| Univariate | 2.331 | 1.037-5.242 | **0.0406** | 2.439 | 0.993-6.376 | 0.0691 |
| Adjusted for clinical variables<sup>c</sup> | 3.230 | 1.268-8.228 | **0.0140** | 3.393 | 0.962-11.97 | 0.0575 |

In the COEUR validation cohort (Table 2a), high epithelial nuclear 53BP1 was significantly associated with poorer disease-specific survival in univariate analysis as a dichotomous variable (hazard ratio (HR) = 2.450; 95% CI: 1.508-3.981; p < 0.001). Statistical significance of this association was maintained in multivariate analysis (HR = 2.325; 95% CI: 1.305-4.141; p= 0.0042). Similar results were obtained for progression-free survival in which high epithelial nuclear 53BP1 was associated with poorer progression-free survival as a dichotomous variable in univariate (HR = 2.651; 95% CI: 1.747-4.023; p < 0.001) and multivariate analyses (HR = 2.179; 95% CI: 1.349-3.518; p = 0.0015). However, these associations were also present in the analysis of epithelial nuclear 53BP1 as a continuous variable, lending support to a causal link between 53BP1 and clinical outcomes. Higher 53BP1 levels were associated with poorer disease-specific survival in univariate (HR = 1.007; 95% CI: 1.002-1.013; p = 0.0134) and multivariate analyses (HR = 1.007; 95% CI: 1.001-1.014; p = 0.0310) as well as with poorer progression-free survival in univariate (HR = 1.010; 95% CI: 1.005-1.016; p < 0.001) and multivariate analyses (HR = 1.009; 95% CI: 1.002-1.015; p = 0.0060).

Similar trends were observed in the CHUM cohort (Table 2b). When epithelial nuclear 53BP1 was analyzed as a dichotomous variable, high levels were significantly associated with disease-specific survival in univariate (HR = 2.331; 95% CI: 1.037-5.242; p = 0.0406) and multivariate analyses (HR = 3.230; 95% CI: 1.268-8.228; p = 0.0140). Non-statistically significant trends towards poorer progression-free survival were observed in univariate (HR = 2.439; 95% CI: 0.993-6.376; p = 0.0691) and multivariate analyses (HR = 3.393; 95% CI: 0.962-11.97; p = 0.0575). Likely due to the smaller number of patients in the CHUM cohort, analysis of 53BP1 as a continuous variable did not reveal any statistically significant associations, but non-significant trends towards poorer outcomes with higher levels of 53BP1 were again observed. Taken together, these results suggest that tissue epithelial nuclear 53BP1 predicts clinical outcomes in treatment-naïve HGSOC patients, independent of other established prognostic variables, and suggests a causal link between 53BP1 and clinical outcomes.

### High *TP53BP1* mRNA levels in whole tumour correlate with poor overall survival

To further validate the relationship between 53BP1 and clinical outcomes, we studied a third cohort. We obtained whole-tumour *TP53BP1* mRNA expression data and the associated clinical data for the Cancer Genome Atlas (TCGA) Firehose Legacy cohort, which included 591 HGSOC patients. The clinical characteristics of patients in the TCGA cohort are presented in Table 3. Given that disease-specific or progression-free survival data were not available, overall survival was used as the clinical end point. Patients were separated into high and low *TP53BP1* mRNA groups as described previously. In univariate analysis (Kaplan-Meier and univariate Cox regression analyses), no significant associations were observed between whole-tumour *TP53BP1* mRNA and overall survival. However, higher *TP53BP1* mRNA levels did trend towards poorer overall survival in Kaplan-Meier (Fig. 3) as well as Cox regression analyses as both dichotomous and continuous variables (Table 4). Remarkably, multivariate Cox regression analysis controlling for age at diagnosis, disease-stage at presentation and residual disease after surgery, revealed that higher levels of *TP53BP1* were significantly associated with poorer overall survival whether analyzed as a dichotomous (HR = 1.206; 95% CI: 1.026-1.416; p = 0.0228) or continuous variable (HR = 1.047; 95% CI: 1.008-1.089; p = 0.0192) (Table 4), further supporting a causal link between 53BP1 and clinical outcomes. Contrary to the COEUR and CHUM cohorts, age at diagnosis in addition to the other two clinical variables that were controlled for were strongly associated with the clinical outcome studied (Table 4). Thus, the absence of an association between *TP53BP1* mRNA and overall survival in univariate analysis is likely due to the lack of censorship of non-disease-specific deaths in the reported clinical outcome. Nonetheless, these data further support the association between 53BP1 and poor clinical outcomes.

**Figure 3:**
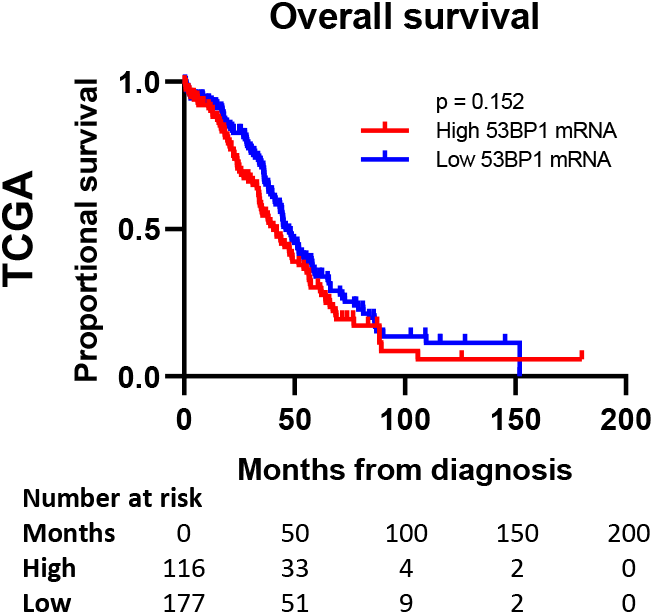
53BP1 mRNA levels trend towards an association with overall survival in the TCGA Firehose Legacy cohort. Kaplan-Meier curves for overall survival in the TCGA Firehose Legacy cohort for high and low expressors of whole tumoral *TP53BP1* mRNA. Statistical significance was calculated using the log-rank test.

**Table 3:** Clinical characteristics of the patients in the TCGA Firehose Legacy cohort.

| Characteristics | TCGA (n = 591) |
| --- | --- |
| Age at diagnosis – years |  |
| Median | 59.0 |
| Range | 26.0-89.0 |
| FIGO stage at presentation – n (%) |  |
| 1 | 17 (2.9) |
| 2 | 29 (4.9) |
| 3 | 441 (74.6) |
| 4 | 89 (15.1) |
| Unknown | 15 (2.5) |
| Residual disease after surgery – n (%) |  |
| None | 119 (20.1) |
| 1 cm or less | 254 (43.0) |
| 1-2 cm | 36 (6.1) |
| Greater than 2 cm | 105 (17.8) |
| Milliary | 0 (0.0) |
| Unknown | 77 (13.0) |
| BRCA-status – n (%) |  |
| BRCA1-mutated | 20 (3.4) |
| BRCA2-mutated | 22 (3.7) |
| Wild-type | 269 (45.5) |
| Unknown | 280 (47.4) |
| First-line chemotherapy – n (%) |  |
| Carboplatin and taxane | 234 (39.6) |
| Cisplatin and taxane | 42 (7.1) |
| Carboplatin | 34 (5.8) |
| Cisplatin | 5 (0.8) |
| Other | 206 (34.9) |
| None | 0 (0.0) |
| Unknown | 70 (11.8) |
| Follow up – months |  |
| Median | 33.1 |
| Range | 0.3-180.1 |
| Overall survival status – n (%) |  |
| Alive | 236 (39.9) |
| Deceased | 337 (57.0) |
| Unknown | 18 (3.0) |
| Overall survival time – months |  |
| Median | 44.49 |
| Range | 0.3-180.1 |
| Tissue year of collection |  |
| Median | 2004 |
| Range | 1992-2013 |

**Table 4:** Cox regression analysis of *TP53BP1* mRNA in the TCGA Firehose Legacy cohort. Cox proportional hazard regression analysis of selected variables and overall survival in the TCGA Firehose Legacy cohort. Age at diagnosis and whole tumoral *TP53BP1* mRNA expression (where stated) were continuous variables. ^a^Hazard ratio (HR) estimated from the Cox proportional hazard regression model. ^b^95% confidence interval around the estimated HR. ^c^Multivariate analysis was adjusted for three clinical variables: age at diagnosis, disease stage at presentation, and residual disease after surgery. Statistically significant p-values in bold.

| Variable | TCGA |  |  |
| --- | --- | --- | --- |
|  | Overall survival |  |  |
|  | HR<sup>a</sup> | 95 % CI<sup>b</sup> | p-value |
| Age at diagnosis | 1.024 | 1.014-1.034 | **2x10<sup>-6</sup>** |
| Disease stage |  |  | **0.00715** |
| 2 vs 1 | 0.991 | 0.256-3.836 | 0.990 |
| 3 vs 1 | 2.330 | 0.746-7.273 | 0.145 |
| 4 vs 1 | 3.074 | 0.967-9.776 | 0.0571 |
| Residual disease after surgery |  |  | **1x10<sup>-6</sup>** |
| 1 cm or less vs none | 2.080 | 1.478-2.926 | **2.6x10<sup>-5</sup>** |
| 1-2 cm vs none | 2.574 | 1.554-4.265 | **2.4x10<sup>-4</sup>** |
| Greater than 2 cm vs none | 2.860 | 1.946-4.202 | **8.7x10<sup>-8</sup>** |
| Miliary vs none | - | - | - |
| **53BP1 mRNA (continuous)** |  |  |  |
| Univariate | 1.025 | 0.987-1.064 | 0.199 |
| Adjusted for clinical variables<sup>c</sup> | 1.047 | 1.008-1.089 | **0.0192** |
| **53BP1 mRNA (high vs. low)** |  |  |  |
| Univariate | 1.242 | 0.923-1.671 | 0.153 |
| Adjusted for clinical variables<sup>c</sup> | 1.206 | 1.026-1.416 | **0.0228** |

### 53BP1 mediates sensitivity to chemotherapy in HGSOC cell lines

Although the relationship between 53BP1 and clinical outcomes was relatively consistent across three cohorts of HGSOC patients, how 53BP1 was tied to clinical outcomes remained unknown. To further explore this question, we opted to probe 10 treatment-naïve cell lines in a panel of well-characterized HGSOC cell lines derived at the CHUM from either ascites or tumour tissue of patients diagnosed with HGSOC (Fleury et al., 2015; Letourneau et al., 2012; Ouellet et al., 2008). Given its extensive involvement in the DDR, we hypothesized 53BP1 must modulate sensitivity to anticancer treatments at a cellular level. Using western blot analysis, we measured native 53BP1 protein levels in the treatment-naïve HGSOC cell lines, as well as in four normal epithelial ovarian cell cultures (Fig. 4a-b). Interestingly, 53BP1 levels in HGSOC cell lines trended higher than in their normal ovarian epithelium counterparts (Fig. 4c), though levels varied significantly among the different cell lines. Nonetheless, this finding was suggestive of a dependence of HGSOC cells on 53BP1 for survival. Next, we plotted 53BP1 protein levels against each cell lines’ sensitivity to carboplatin, a first-line drug for the treatment for HGSOC. The IC_50_ for a 24-hour treatment with this drug on colony formation assay was previously reported in eight of the 10 cell lines (Brodeur et al., 2021; Fleury et al., 2015; Letourneau et al., 2012); it was determined for this study for the remaining two cell lines (Supplemental Table 1). Carboplatin IC_50_ values were plotted against the 53BP1 protein levels we had determined for each cell line and revealed a striking correlation (Pearson r = 0.6783; p = 0.0311) (Fig. 4d), which showed that cell lines expressing higher levels of 53BP1 had a lower sensitivity to carboplatin.

**Figure 4:**
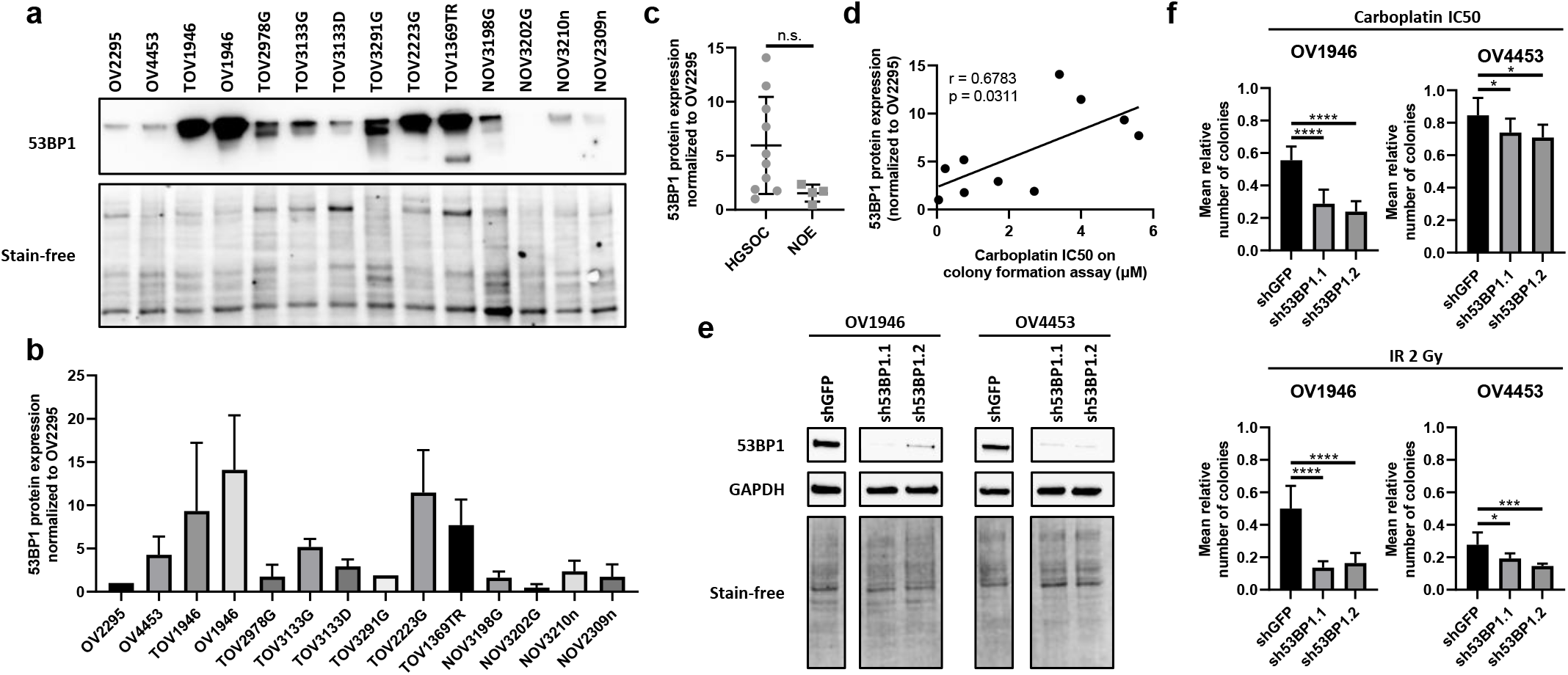
53BP1 mediates sensitivity to chemotherapy in HGSOC cell lines. (**a**) Representative western blot for 53BP1 in 10 treatment-naïve HGSOC cell lines and four normal ovarian epithelium cultures (last four lanes)? as well as the associated stain-free loading control. The western blot was repeated three times. (**b**) Quantification of 53BP1 from the western blot in (**a**). The 53BP1 chemiluminescence signal was normalized to the stain-free signal in each cell line and was then averaged across three technical replicates. Data is reported as protein levels normalized to OV2295. (**c**) Average 53BP1 protein levels in 10 HGSOC cell lines as compared to those in four normal ovarian epithelium (NOE) cultures. Hypothesis testing was carried out using the Student t-test. n.s. non-significant. (**d**) Pearson correlation between IC_50_ for a 24-hour carboplatin treatment in colony formation assays and 53BP1 protein levels in 10 HGSOC cell lines. Colony formation assays to determine IC_50_ were carried out three times in each cell line. (**e**) Western blot for 53BP1 in OV1946 and OV4453 HGSOC cells expressing shGFP, sh53BP1.1 and sh53BP1.2. These western blots were repeated three times each. (**f**) Colony formation assays with OV1946 and OV4453 HGSOC cells expressing shGFP, sh53BP1.1 and sh53BP1.2 after 24-hour treatment with carboplatin at their previously established IC_50_ (top) or 2 Gy ionizing radiation (IR; bottom). Colony numbers were normalized to their untreated controls. Hypothesis testing was carried out using the Student t-test. *p < 0.05, ***p < 0.001, and ****p < 0.0001. Each colony formation assay was performed in quadruplicate and repeated at least two times.

To firmly establish a causal role for 53BP1 in the cellular sensitivity to DNA damaging anticancer strategies, we performed stable 53BP1 depletion in HGSOC cells and evaluated their subsequent colony forming ability in response to various antineoplastic treatments. One cell line highly expressing 53BP1, OV1946, as well as one weakly expressing 53BP1, OV4453, were modified to express either of two short hairpins RNAs (shRNA) targeting 53BP1 (sh53BP1.1 and sh53BP1.2) or a control shRNA targeting green fluorescent protein (shGFP) (Fig. 4e). Colony formation assays were then performed in three conditions: untreated control, a 24-hour carboplatin treatment at previously determined IC_50_, and 2 Gy ionizing radiation.

Knockdown of 53BP1 significantly sensitized both the high and the low native 53BP1 cell lines to carboplatin. Unsurprisingly, the strength of the effect was markedly less in the cell line with low native 53BP1. Both cell lines were also sensitized to 2 Gy ionizing radiation (Fig. 4f), which induces different types of DNA damage, and subsequent DNA repair from those involved in carboplatin toxicity (Basu and Krishnamurthy, 2010; Hur and Yoon, 2017). Altogether, these results suggest that 53BP1 mediates sensitivity to DNA-damaging agents, including first-line chemotherapy, in HGSOC at a cellular level, and this may explain the association of 53BP1 and HGSOC treatment outcomes at the patient level.

## DISCUSSION

In this study, we show that high nuclear 53BP1 protein levels in tumour tissue were associated with poorer disease-specific survival and progression-free survival in two independent HGSOC cohorts, and that high whole-tumour *TP53BP1* mRNA levels were also associated with poorer overall survival in a third cohort. Remarkably, these associations persisted or became statistically significant when we controlled for established prognostic factors that included age at diagnosis, disease stage at presentation and residual disease after surgery, suggesting that 53BP1 may be an independent prognostic marker in treatment-naïve patients. To our knowledge, this is the first extensive study on how 53BP1 relates to patient outcomes in HGSOC based on a large number of patients from three independent cohorts that includes both quantitative protein and mRNA expression data.

Consistent with our observations, Pennington et al. were among the first to report a potential link between poor prognosis and high 53BP1 in epithelial ovarian cancer. They showed that low tumoral *TP53BP1* mRNA expression was associated with improved overall survival in wild type *BRCA1* tumours in an 89-patient subgroup of their cohort (Pennington et al., 2013). However, they found no such association between the frequency of 53BP1-positive tumor cells as evaluated by immunohistochemistry in the entire cohort of 248 patients. Intriguingly, in this ovarian cancer cohort *BRCA1* mutations were associated with higher levels of 53BP1 protein in tumour tissues (Pennington et al., 2013). However, we found no association between 53BP1 and *BRCA1* or *BRCA2* mutations in either the COEUR validation or CHUM HGSOC cohorts (Supplemental Fig. 1). To probe the link between 53BP1 and treatment response in a different manner, Hurley et al. found in a small cohort of ovarian cancer patients with HR deficiency that low tumoral 53BP1 determined using immunohistochemistry was associated with a weak short-term tumoral response to PARP inhibition(Hurley et al., 2019). This is consistent with the reported role of 53BP1 in DNA repair pathway choice and the restoration of HR in *BRCA1*-deficient cells following the downregulation of 53BP1 (Bunting et al., 2010; Chapman et al., 2013). Measured parameters and patient cohorts differ substancially between our study and the previous ones, but taken together they suggest the possibility that 53BP1 levels differentially impact ovarian tumors depending on their HR status, or perhaps 53BP1 expression may reflect BRCA status in HGSOC, as is the case in breast cancer (Jacot et al., 2013).

How the high expression of 53BP1 is tied to poor outcomes in HGSOC remains unclear, but several of our observations suggest a direct causal relationship. In this study, we found that 53BP1 as a continuous variable was associated with poor clinical outcomes in two of the cohorts we studied. Additionally, we observed that native 53BP1 levels in HGSOC cell lines correlated strikingly well with their sensitivity to carboplatin. Lastly, knockdown of 53BP1 in two HGSOC cell lines increased their sensitivity to carboplatin, a first-line chemotherapy agent. Thus, 53BP1 may directly mediate treatment outcomes, possibly by modulating cellular responses to cytotoxic chemotherapy. Our cell culture results offer this mechanistic explanation for the results obtained with the clinical data. A rescue experiment for the knockdown of 53BP1 was not performed due to the technical difficulty posed by the large size of the *TP53BP1* gene. Nonetheless, the use of two separate shRNAs targeting *TP53BP1* and of a control shRNA minimizes the risk of results generated from off-target effects.

Despite demonstrating a strong association between 53BP1 and clinical outcomes in HGSOC, our results do not allow us to further clarify the underlying molecular mechanisms. 53BP1 is involved in the DDR at multiple levels (Mirman and de Lange, 2020), which includes acting as a scaffold for several DDR factors at sites of DNA damage (Gong et al., 2009; Huen et al., 2010; Silverman et al., 2004) and promoting NHEJ for the repair of DNA DSBs (Panier and Boulton, 2014). However, it is unlikely that 53BP1 promotes rapid repair of DNA damage resulting from cytotoxic chemotherapies as the knockdown of 53BP1 has been shown to only minimally slow down DNA repair (Noon et al., 2010). More recently it has been reported that the interaction between 53BP1 and p53 may modulate the transactivation function of p53, and thus, cell fate decisions downstream of the DDR (Cuella-Martin et al., 2016). Although the near ubiquitous mutation of p53 in HGSOC (Cancer Genome Atlas Research, 2011) potentially renders this explanation unlikely as well, it cannot be excluded that higher levels of 53BP1 may potentiate the oncogenic functions of mutant p53 in HGSOC (Zhang et al., 2016). On a different note, several groups have shown that 53BP1 is involved in the reinforcement of the S and G2-M checkpoints, particularly in response to low levels of DNA damage (DiTullio et al., 2002; Fernandez-Capetillo et al., 2002; Wang et al., 2002; Ward et al., 2003), and that this checkpoint reinforcement may be mediated by the promotion of phosphorylation of the kinases ATM and CHK2 (DiTullio et al., 2002; Fernandez-Capetillo et al., 2002; Ward et al., 2003). Accordingly, knockdown of 53BP1 in breast cancer cells leads to a decrease of their accumulation in G2-M (Li et al., 2013). Thus, in HGSOC, 53BP1 may help enforce remaining cell cycle checkpoints in the absence of a functional p53 to prevent the progression of cell division in the presence of DNA damage and to maintain some genomic stability. In keeping with this idea, a knockout of 53BP1 in mice with a p53 knockout resulted in the development of lymphomas characterized by signs of genomic instability (Morales et al., 2006). Interestingly, our lab has recently shown that HGSOC cells may enter a senescence-like state to escape PARP inhibitor toxicity, and entry into this senescence-like state involves activation of CHK2 (Fleury et al., 2019), but it was not studied whether 53BP1 was involved in this phenomenon.

Studies examining the prognostic value of 53BP1 in other cancers do not show a consistent association with clinical outcomes. High 53BP1 was associated with poor outcomes in colorectal cancer (Bi et al., 2015) and with aggressive pathological features in papillary thyroid carcinoma (Mussazhanova et al., 2013), while low 53BP1 was associated with a poorer prognosis in breast cancer (De Gregoriis et al., 2017; Jacot et al., 2013; Schouten et al., 2016). Reports are similarly inconsistent at a cellular level. Knockdown of 53BP1 was found to decrease the sensitivity of breast cancer cells to 5-FU (Li et al., 2013) and to decrease the sensitivity of colorectal cancer cells to ionizing radiation (Xiao et al., 2016), whereas it increased the sensitivity of laryngeal cancer cells to ionizing radiation (Gou et al., 2015). This lack of consistency in the literature across different cancer types likely reflects the involvement of 53BP1 at multiple levels in the DDR and suggests that its net effect depends on the integrity of the multiple pathways that interact with 53BP1.

In summary, our findings in three independent cohorts suggest that 53BP1 is associated with clinical outcomes to current first-line chemotherapies in treatment-naïve HGSOC patients, independent of established prognostic factors. This association may be the result of 53BP1-mediated variations in sensitivity to chemotherapeutic treatments at a cellular level, suggesting that 53BP1 modulation could represent a novel treatment target for HGSOC.

## MATERIALS AND METHODS

### Patients and tissues

HGSOC TMAs were constructed from formalin-fixed, paraffin-embedded (FFPE) tissues that were obtained from the Terry Fox Research Institute’s Canadian Ovarian Exploratory United Resource (COEUR) and from the CHUM. FFPE tissues were collected during primary cytoreductive surgery of patients with HGSOC. Patient consent was obtained prior to sample collection. Patients from each biobank were included in the study if review by a pathologist confirmed HGSOC histology and if they had not received neoadjuvant chemotherapy prior to cytoreductive surgery.

The COEUR validation cohort included tissues from 173 patients, recruited between 1992 and 2011, at 10 different Canadian biobanks, including the CHUM. These patients constitute a subset of the complete TFRI-COEUR cohort previously described (Le Page et al., 2018). Each tumour was represented by duplicate tissue cores. Histopathology was evaluated by pathologists specializing in gynecological oncology (Kurosh Rahimi and Martin Köbel). The clinical characteristics of the patients included in the COEUR validation cohort are presented in Table 1.

The CHUM cohort included tissues from 76 patients recruited between 1993 and 2012. Patients represented in both the CHUM cohort and COEUR validation cohort were excluded from the CHUM cohort and instead included in the COEUR validation cohort for analysis (n = 20; the final number of patients included in the CHUM cohort for analysis was 56). Each tumour was represented by duplicate tissue cores. Histopathology was evaluated by a pathologist specializing in gynecological oncology (Kurosh Rahimi). The disease stage was determined at the time of surgery according to the criteria established by the International Federation of Gynecology and Obstetrics. Likewise, residual disease was determined at the time of surgery. The clinical characteristics of the patients included in the CHUM cohort are shown in Table 1.

### Study approval and ethics statement

Approval from the CHUM institutional ethics committee was obtained. Informed consent from all patients was obtained before sample collection and samples were anonymized.

### The Cancer Genome Atlas (TCGA) Firehose Legacy dataset

Whole-tumour *TP53BP1* mRNA expression data was obtained for 591 HGSOC tumour samples from the TCGA Firehose Legacy Ovarian Serous Cystadenocarcinoma dataset (doi:10.7908/C11G0KM9) in January 2017. Associated clinical data was obtained at the same time. All samples were included in the analysis as all were considered to have HGSOC histology and none had been treated with neoadjuvant chemotherapy prior to sample collection. Contrary to the COEUR validation and CHUM cohorts, the outcome of interest for the TCGA dataset was overall survival as data for disease-specific survival and progression-free survival were unavailable. Select clinical characteristics for patients included in the TCGA dataset are presented in Table 3.

### Antibodies

Details of primary and secondary antibodies used in this study are summarized in Supplemental Table 2.

### Tissue immunofluorescence (IF) staining

The TMA block was sectioned into 4-µm thick slices and processed using a Benchmark XT automated stainer (Ventana Medical System Inc., Tucson, AZ, USA). Antigen retrieval was carried out with Cell Conditioning 1 (Ventana Medical System Inc.; no. 950-124) for 60 minutes. The prediluted primary antibody was automatically dispensed and incubated for 60 minutes at 37°C. The following steps were performed manually. After blocking with Protein Block serum-free reagent (Dako #X0909), secondary antibodies were added for 45 minutes, followed by washing and blocking overnight at 4°C with diluted Mouse-On-Mouse reagent (Vector #MKB-2213). This was followed by an incubation of 60 minutes with primary antibodies of the epithelial mask. The Alexa Fluor 750 secondary antibody was incubated for 45 minutes, followed by DAPI staining for 5 minutes. Staining with 0.1% Sudan black for 15 minutes to quench tissue autofluorescence was followed by coverslip mounting using Fluoromount Aqueous Mounting Medium (#F4680, Sigma). The full TMA was scanned-digitalized using a 20X 0.75NA objective with a resolution of 0.325 μm (BX61VS, Olympus). Multicolor images were segmented and quantified using Visiopharm Integrator System (VIS) version 4.6.1.630 (Visiopharm, Denmark).

### VIS analysis

IF scoring was performed using the VIS software. Image segmentation is summarized in Figure 1b. Briefly, VIS was used to first identify the cores by separating them from the background. They were then separated into epithelial and stromal compartments according to the intensity of the cytokeratin 7, 18 and 19 staining. Nuclei were then identified according to the intensity of the DAPI signal. Cores were then separated into regions of interest according to these compartments. VIS was then used to measure the MFI of all pixels in the original digitalized multicolor TMA images for a chosen region of the tissue core (i.e., the epithelial nuclei). When replicate cores were available for a single patient, the MFIs measured for each core were averaged in order to give a single average MFI value for each patient. Reproducibility between duplicate cores is presented in Supplemental Figure 2.

### Statistical analysis for the TMA

Patients were included in the statistical analysis only if they satisfied both of the following criteria: 1) their disease had been pathologically classified as HGSOC, and 2) they had not received chemotherapy for ovarian cancer prior to collection of the tissue sample. Statistical analysis was performed using IBM SPSS Statistics for Windows, version 25 (IBM Corp., Armonk, N.Y., USA). Statistical significance was set at p ˂ 0.05. Patients were separated into high or low levels of epithelial nuclear 53BP1 based on the optimal cut-off point for the average MFI of epithelial nuclear 53BP1 found on the receiver operating characteristic curve for each outcome evaluated: disease-specific survival and progression-free survival. For the TCGA Firehose Legacy cohort, patients were separated into high or low *TP53BP1* mRNA expressors based on the optimal cut-off point for whole-tumour *TB53BP1* mRNA found on the receiver operating characteristic curve for overall survival. Kaplan-Meier survival analysis was performed for disease-specific survival and progression-free survival for the COEUR validation and CHUM cohorts, and for overall survival for the TCGA Firehose Legacy cohort. Statistical significance was calculated using the log-rank test. Cox regression analysis was performed using epithelial nuclear 53BP1 levels (or whole-tumour *TP53BP1* mRNA levels) as a dichotomous variable (with high and low groups previously determined as described above) as well as a continuous variable using the average MFI values for each patient (or the *TP53BP1* mRNA level). Univariate and multivariate analyses were performed. Multivariate analysis was performed with adjustment for age at diagnosis (entered as a continuous variable), disease stage at presentation (entered as a categorical variable), and residual disease after surgery (entered as a categorical variable).

### Cell lines and cell culture

Ten human HGSOC cell lines used in this study: OV2295, OV4453, TOV1946, OV1946, TOV2978G, TOV3133G, TOV3133D, TOV3291G, TOV2223G, and TOV1369TR. They were derived in Dr. Mes-Masson’s laboratory from the ascites or tumour tissues of patients diagnosed with HGSOC and have been extensively characterized (Fleury et al., 2015; Letourneau et al., 2012; Ouellet et al., 2008). Four different normal epithelial ovarian cell lines established in Dr. Mes-Masson’s laboratory were also used: NOV3198G, NOV3202G, NOV3201n and NOV2309n. All cell lines were maintained in a low oxygen condition (7% O_2_ and 5% CO_2_) and grown in OSE medium (Wisent, Montreal, QC) with 10% fetal bovine serum (FBS) (Wisent), 0.5 μg/ml amphotericin B (Wisent) and 50 μg/ml gentamicin (Life Technologies Inc., Burlington, ON).

### Cloning, viruses and infections

Viruses were produced as described previously (Campeau et al Plos One 2009, Rodier NCB 2009). Cells were infected with virus titrated to achieve 90% infectivity with 4 μg/mL of polybrene in a final volume of 1 mL in 6-well plates. Infections were followed 48 hours later by puromycin selection (1 µg/mL).

The shRNAs lentivectors against 53BP1 and shGFP (green fluorescent protein, control) were purchased from Open Biosystems (Horizon discovery) (lentiviral PLKO.1 vector with puromycin selection). The following sequences were used for each shRNA:

sh53BP1.1: 5’-AAACCAGTAAGACCAAGTATC -3’;

sh53BP1.2: 5’-AATCAATACTAATCACACTGG -3’;

shGFP: 5’-GCTGGAGTACAACTACAAC-3’.

### Protein preparation and western blot analysis

Whole cell lysates were prepared by scraping cells with mammalian protein extraction reagent (MPER, Thermo Fisher Scientific, Waltham, MA) containing a protease and phosphatase inhibitor cocktail (Sigma-Aldrich Inc., St. Louis, MO). Protein concentration was measured using the bicinchoninic acid (BCA) protein assay (Thermo Fisher Scientific). Ten micrograms of total protein extract underwent electrophoresis in stain-free 4–15% gradient Tris-glycine SDS-polyacrylamide gels (Mini PROTEAN® TGX Stain-Free™ Gels, Bio-Rad Laboratories, Hercules, CA) and transferred onto PVDF membranes (Hybond-C Extra, GE Healthcare Life Sciences, Mississauga, ON, Canada). Membranes were blocked with 5% bovine serum albumin (BSA) in PBS for 1 hour and probed with primary antibodies overnight at 4 °C. Bound primary antibodies were detected with peroxidase-conjugated secondary antibodies (Cell Signaling Technology Inc., Danvers, MA) and enhanced chemiluminescence (Thermo Fisher Scientific). Chemiluminescence was detected using the ChemiDoc MP Imaging System (Bio-Rad Laboratories). Protein loading control was evaluated using the stain-free technology (Bio-Rad Laboratories) and/or GAPDH. The western blot was carried out three separate times to assess native levels of 53BP1 in HGSOC cell lines and to assess the efficacy of the shRNAs. Quantification of the protein bands were performed using Image Lab 6.1 Software (Bio Rad). Quantification of the 53BP1 bands was normalized using the total protein quantity detected by the stain-free images.

### Carboplatin IC_50_ determination by clonogenic assay

Carboplatin IC_50_ determination in HGSOC cell lines has previously been described (Fleury et al., 2015). IC_50_ was assessed by colony formation assay. Cells were seeded in a 6-well plate at the appropriate density for each cell line to allow formation of individual colonies. After cells were allowed to adhere for 16 hours, the media were removed and replaced with OSE medium prepared as above and containing 0-100 µM of carboplatin (Hospira Healthcare Corporation, Saint-Laurent, QC). Cells were incubated with the treatment for 24 hours. The carboplatin-containing media were removed and replaced with carboplatin-free media. Cells were incubated until colonies became visible at a 2X magnification. Cells were then fixed using cold methanol and stained with a mix of 50 % v/v methanol and 0.5 % m/v methylene blue (Sigma-Aldrich Inc., St. Louis, MO). Colonies were counted under a stereomicroscope. The number of counted colonies was then normalized to the control condition. IC_50_ values were determined using Graph Pad Prism 5 software (GraphPad Software Inc., San Diego, CA). Each individual experiment was performed in triplicate and repeated three times.

### Correlation for 53BP1 and carboplatin IC_50_

Mean normalized 53BP1 protein levels in each cell line obtained on western blot were plotted against measured carboplatin IC_50_ values obtained by colony formation assay. A Pearson correlation coefficient and the associated p-value were calculated using Graph Pad Prism 8 software (GraphPad Software Inc., San Diego, CA).

### Clonogenic assays with chemotherapy and ionizing radiation

OV1946 and OV4453 cells expressing either shGFP, sh53BP1.1 or sh53BP1.2 were seeded in 6-well plates at the appropriate density for each cell line to allow formation of individual colonies. After cells were allowed to adhere for 16 hours, the media were removed and replaced with OSE medium prepared as above and containing either 3.40 µM (OV1946) or 0.23 µM (OV4453) of carboplatin (Hospira Healthcare Corporation, Saint-Laurent, QC) for the carboplatin condition, or OSE medium prepared as above without carboplatin for the 2 Gy ionizing radiation condition and control condition. Cells for the ionizing radiation condition were subjected at this time to 2 Gy ionizing radiation. Cells were then incubated with their respective media for 24 hours. The carboplatin-containing media were removed and replaced with carboplatin-free media, whereas media for conditions without carboplatin were simply changed for more carboplatin-free media. Cells were incubated until colonies became visible at a 2X magnification. Cells were fixed using cold methanol and stained with a mix of 50 % v/v methanol and 0.5 % m/v methylene blue (Sigma-Aldrich Inc., St. Louis, MO). Colonies were counted under a stereomicroscope. The number of counted colonies was normalized to the control condition. Each individual experiment was performed with four technical replicates and repeated at least twice.

## ACKNOWLEDGEMENTS

We thank members of the Mes-Masson, Provencher and Rodier laboratory for valuable comments and discussions. We thank the CRCHUM molecular pathology core facility and the Institut du cancer de Montréal (ICM) Imaging and Live imaging platform. This work was supported by the ICM (DP, AMMM, FR) and by the Canadian Institute for Health Research (CIHR MOP114962 & CIHR PJT173341 to FR), the Terry Fox Research Institute (TFRI 1030 to FR) and the Cancer Research Society (CRS) in partnership with Ovarian Cancer Canada (20087 to AMMM, DP; 22713 to FR). AMMM, DP and FR are researchers of the CRCHUM/ICM, which receive support from the Fonds de recherche du Québec - Santé (FRQS). FR is supported by a FRQS junior I-II-senior career awards (22624, 33070, 281706). Ovarian tumor banking was supported by the Banque de tissus et de données of the Réseau de recherche sur le cancer of the FRQS affiliated with the Canadian Tumor Repository Network (CTRNet). LCG and MS received Canderel fellowships from the ICM, SC received a FRQS postdoctoral award, MS received a CIHR doctoral award and YZ was supported by an ICM/MITACS postdoctoral fellowship. This study uses resources provided by the COEUR biobank funded by the TFRI and managed and supervised by the CHUM. The Consortium acknowledges contributions to its COEUR biobank from Institutions across Canada (for a full list see www.tfri.ca/COEUR/members).

## AUTHORS CONTRIBUTIONS

MS and FR designed the study. MS, JB, NM, HF, LC and IC performed experiments. KR revised all pathology samples for TMA construction. MS, JB, and FR collected and analyzed data. MS and FR wrote the manuscript. AMMM and DP provided technical support, biobank access, molecular pathology expertise, conceptual advice, and revised the manuscript.

