## Supplemental figures for "53BP1 mediates sensitivity to chemotherapy and is associated with poor clinical outcomes in high-grade serous ovarian cancer"

**Supplemental Table 1: Carboplatin $IC_{50}$ of HGSOC cell lines as determined by colony formation assay.**

| Cell line | Carboplatin $IC_{50}$ ± SEM (μM) |
|---|---|
| OV2295 | 0.05 ± 0.01 |
| OV4453 | 0.23 ± 0.074 |
| TOV1946 | 5.2 ± 1.1 |
| OV1946 | 3.4 ± 0.18 |
| TOV2978G | 0.76 ± 0.24 |
| TOV3133G | 0.75 ± 0.63 |
| TOV3133D | 1.7 ± 0.99 |
| TOV3291G | 2.7 ± 0.9 |
| TOV2223G | 4.0 ± 0.7 |
| TOV1369TR | 5.6 ± 1.3 |

**Supplemental Figure 1: 53BP1 expression as a function of BRCA status in the COEUR and CHUM cohorts.**

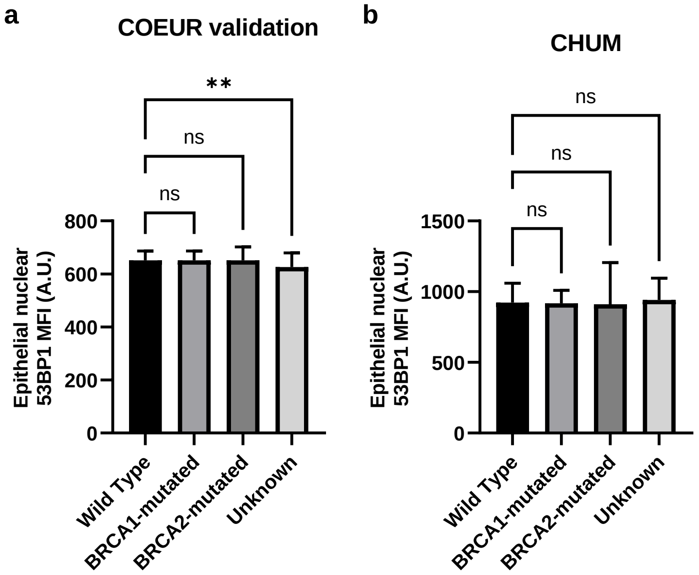

**Supplemental Table 2: Summary of primary and secondary antibodies used in this study.**

| Antibody (clone, if monoclonal) | Species | Manufacturer (catalog number) | Dilution for TMA IF | Dilution for western blot |
|---|---|---|---|---|
| **Primary antibodies** | | | | |
| 53BP1 | Rabbit | Novus Biologicals (NB100-305) | 1:500 | 1:2000 |
| GAPDH | Rabbit | Cell Signaling (2118S) | - | 1:2000 |
| **Epithelial mask primary antibodies** | | | | |
| Cytokeratin 7 (OV-TL 12/30) | Mouse | ThermoFisher Scientific (MS-1352-P) | 1:200 | - |
| Cytokeratin 18 (DC-10) | Mouse | Santa-Cruz (sc-6259) | 1:200 | - |
| Keratin 19 (A53-B/A2.26) | Mouse | ThermoFisher Scientific (MS-198-P) | 1:200 | - |
| **Secondary antibodies** | | | | |
| Alexa-Fluor 647 anti-rabbit | Donkey | Life Technologies (A31573) | 1:250 | - |
| Alexa-Fluor 750 anti-mouse | Goat | Life Technologies (A21037) | 1:250 | - |
| Anti-rabbit IgG, HRP-linked | Goat | Cell Signaling (7074P2) | - | 1:5000 |

**Supplemental Figure 2: Correlations of mean fluorescent intensity of 53BP1 between duplicate cores in the COEUR and CHUM cohorts.**

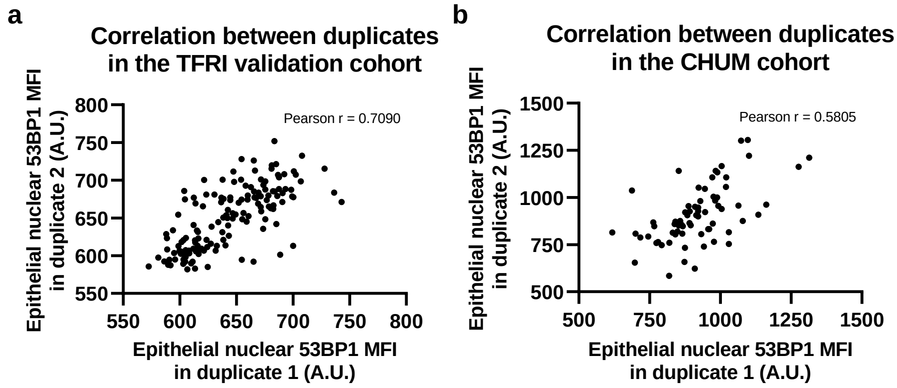
